# MutSigCVsyn: Identification of Thirty Synonymous Cancer Drivers

**DOI:** 10.1101/2022.01.16.476507

**Authors:** Yiyun Rao, Nabeel Ahmed, Justin Pritchard, Edward O’Brien

**Affiliations:** Huck Institute of the Life Sciences, Pennsylvania State University, University Park, 16802, Pennsylvania, USA; Department of Biomedical Engineering, Pennsylvania State University, University Park, 16802, Pennsylvania, USA; Department of Chemistry, Pennsylvania State University, University Park, 16802, Pennsylvania, USA; Institute for Computational and Data Sciences, Pennsylvania State University, University Park, 16802, Pennsylvania, USA

## Abstract

Synonymous mutations, which change only the DNA sequence but not the encoded protein sequence, can affect protein structure and function, mRNA maturation, and mRNA half-lives. The possibility that synonymous mutations can act as cancer drivers has been explored in several recent studies. However, none of these studies control for all three levels (patient, histology, and gene) of mutational heterogeneity that are known to affect the accurate identification of non-synonymous cancer drivers. Here, we create an algorithm, MutSigCVsyn, an adaptation of MutSigCV, to identify synonymous cancer drivers based on a novel non-coding background model that takes into account the mutational heterogeneity across these levels. Examining 2,572 PCAWG cancer whole-genome sequences, MutSigCVsyn identifies 30 novel synonymous drivers that include mutations in promising candidates like BCL-2. By bringing the best practices in non-synonymous driver identification to the analysis of synonymous drivers, these are promising candidates for future experimental study.

## Introduction

‘Driver’ mutagenic events confer a selective growth advantage to cells and contribute to tumorigenesis^1,2^. Discovering and characterizing these cancer driver genes using large-scale cancer genome sequencing data is a major component of modern cancer research^2,3^. These drivers are typically identified through aberrantly high mutation rates in specific genes relative to an estimate of the background mutation rate^4–6^. Classic efforts have identified a “long-tail” distribution of cancer driver mutations, where some mutations (*e.g*., KRAS G12D^7^) are highly prevalent, and other mutations are extraordinarily rare^8,9^. However, many tumors do not harbor any known cancer drivers. A reasonable assumption is that these tumors harbor driver mutations that are rare enough to be undetectable in existing cohorts^10^. The unambiguous detection of these novel long-tail drivers is a challenge because of the underpowered sample size of many cohorts^11^. However, it may also be a challenge because research labs have primarily looked for cancer drivers involving non-synonymous mutations or non-coding mutations in promoters and other regulatory regions^12,13^.

Synonymous mutations are one class of historically disregarded mutations that might be long-tail drivers. Synonymous mutations alter the mRNA coding sequence but not the encoded protein’s primary structure. In the past, these mutations were assumed to be phenotypically “silent”^14,15^. Nonetheless, synonymous codons encode information beyond amino acids. Protein structure and function can be altered by introducing synonymous mutations that change the rate of protein translation^16–18^. Such variation had been found to affect co-translational folding^19^, translational accuracy^20^, and posttranslational modifications^21^. Additionally, synonymous mutations also play a regulatory role in transcription by altering mRNA structure^22^, and in some cases affecting the mRNA splicing process^23^. Both of these translational and transcriptional effects had been found to impact cell fitness in bacteria^18,24^, and linked to a number of human diseases^25^. It is now broadly accepted that synonymous mutations can affect subcellular processes and phenotype^26,27^.

Two sets of evidence indicate that selective constraints act at synonymous mutation positions in cancer, suggesting a functional role. First, bioinformatic analyses indicate a global selection for synonymous mutations in oncogenes. Supek et al.^28^ found that the synonymous mutation rate is elevated in oncogenes, especially near exon-intron boundaries, regardless of local mutation rates. Analyses from Chu et al.^29^ on single nucleotide polymorphisms (SNPs) in healthy patients suggested synonymous SNP sites in cancer-related genes may undergo a selection constraint, and are more conservative in oncogenes than in other cancer-related genes. In addition, results from Benisty et al.^30^ suggest that the frequently mutated oncogene in oncogene families (e.g., KRAS) may adapt codon usage to promote cancer cell proliferation. Second, circumstantial evidence connects synonymous mutations and cancer. For example, synonymous mutations in the MDR1 gene, which encodes the efflux pump Pgp, contribute to chemotherapy resistance^31^. In cancer cells, synonymous SNPs in MDR1 affect P-glycoprotein substrate specificity. And synonymous mutations in BAP1 were found to cause exon11 skipping, generating a premature stop codon, and thus a complete loss of function for BAP1^32^. These findings suggest that synonymous mutations are possible long-tail drivers.

To identify synonymous cancer drivers, one of the key aspects is the creation of a comprehensive model to estimate background synonymous mutation rates. Several studies have used a variety of computational approaches^33–36^. The background models in these studies have ranged in complexity and sophistication. For example, in the seminal study by Supek et al.^33^, thirteen covariates at the gene level controlled for regional mutation variation between noncancer genes and oncogenes of interest, but patient-level biases were not accounted for. In another study, Sharma et al.^34^ examined and ranked common synonymous mutations in COSMIC^37^ (a curated database of somatic mutations in cancer) and combined this with orthogonal data including mRNA secondary structural change predictions as well as evolutionary conservation score. However, this approach did not have a formal estimate of the background synonymous mutation frequency. No approach to date has accounted for all three levels of patient-, gene- and disease-specific mutational heterogeneity that are known to lead to inaccurate results in non-synonymous cancer identification^4,38^, and are certain to affect the identification of synonymous drivers.

Controlling for patient-, gene- and disease-specific mutation biases is exemplified by the MutSigCV^4^ algorithm that is the community standard for driver identification in non-synonymous mutations. Here, we bring this same level of background mutational modeling to synonymous mutations by creating an algorithm we refer to as MutSigCVsyn, which allows us to detect synonymous drivers while controlling for confounding mutational biases. This approach is enabled by The Pan-Cancer Analysis of Whole Genomes (PCAWG) sequencing data^39^ from which we use the non-coding mutations within genic regions to adjust for triplet nucleotide mutation biases across diverse patients, tumor histologies, and genes. With this approach, we identify 30 novel synonymous candidate drivers across 18 histology cohorts.

## Methods

### Dataset

The patient MAF (Mutation Annotation Format) files, wig coverage files, RNA-seq data, and cancer driver data were retrieved from the PCAWG portal (https://dcc.icgc.org/releases/PCAWG). Driver genes in Cancer Gene Census were retrieved from the COSMIC website (https://cancer.sanger.ac.uk/cosmic). Gene sequence and annotation data were downloaded from Gencode Release19 website (https://www.gencodegenes.org/human/release_19.html). CERES scores were retrieved from the DepMap web portal (https://depmap.org/portal/)

### Patient and Geneset

2572 PCAWG white-list^39^ patients whose SNV mutation information and wig coverage files both exist were selected. The patients were divided into 39 histology cohorts based on the PCAWG annotation. 139 patients who have total mutation number > 50,000^13^ were defined as hypermutators and were excluded from MutSigCVsyn analysis.

Only protein-coding genes were selected for MutSigCVsyn analysis. As the PCAWG SNVs (Single Nucleotide Variants) were annotated based on Gencode v19, known proteincoding genes in Gencode v19 were selected based on filter “KNOWN” and “protein_coding” in the Gencode v19 gene annotation file. The ‘principal’^40^ transcript, if exists, was used. Otherwise, the longest transcript was used. To make sure all mutations were correctly accounted for in the coverage file as in the MAF file, genes of which the coding/intron/UTR SNV positions don’t match between the MAF files and the coverage files were excluded. This left a final gene set of size 18638 for analysis.

### Preprocess of MutSigCVsyn inputs

#### MAF File Preparation

Mutations of PCAWG patients were annotated via customized script (available on GitHub) into 7 mutation categories based upon the mutational context as in MutSigCV^4^. The categories are:

1. transition mutations at CpG dinucleotides
2. transversion mutations at CpG dinucleotides
3. transition mutations at C:G base pairs not in CpG dinucleotides
4. transversion mutations at C:G base pairs not in CpG dinucleotides
5. transition mutations at A:T base pairs
6. transversion mutations at A:T base pairs
7. null and Indel mutations

#### Coverage File Preparation

The coverage for every single patient at every genomic position in the geneset was calculated based on the wig file to ensure accurate coverage, instead of a simple full coverage model. The calculation process was re-engineered as in the original MutSigCV. One covered genomic position was counted as 1. It was equally divided into 3 parts because the nucleotide has 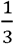 chance to mutate to any of the rest nucleotides (i.e. A could be mutated to C/G/T). Each possible mutation has its consequence, which consists of 3 mutation zones:

1. Synonymous
2. Nonsynonymous
3. Non-coding (Defined as intronic and untranslated regions)

and mutation categories 1 to 6 defined above. The coverage for category 7, the null and indel mutation, was the coverage of the entire gene, which was the sum across categories 1 to 6. These consequences constituted 21 bins in total for each gene. For every position in a gene, the 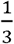 mutation counts were assigned to the corresponding bins and the summed counts were the category-specific coverage for the gene. Full coverage was assumed for unreported positions in the wig file.

#### Covariate file re-annotation

The covariate file provided for MutSigCV^4^ was adopted. However, to avoid the inconsistency of gene naming between the BROAD Institute and PCAWG, the gene names in the covariate file were re-annotated in MutSigCVsyn. All synonyms of the PCAWG gene names were identified using the R package BiomaRt. 862 synonym names were mapped to the BROAD original covariate file and replaced by the new name to generate a new gene covariate file, while the expression, replication timing, and chromatin status data remained the same.

#### Gene dictionary file

MutSigCVsyn only takes mutations in intron and UTR (Untranslated region) into account to avoid transcription-associated mutation bias. Therefore, mutations in the regions that are not transcribed, such as intergenic, promoter, up-/downstream regions, were excluded by removing the variant classification in the gene dictionary file and weren’t recognized in MutSigCVsyn.

### MutSigCVsyn workflow

MutSigCVsyn is adopted from MutSigCV^4^. Several key changes were made to identify the synonymous mutations. A more detailed and technical overview changes can be found in the GitHub repository (https://github.com/ryy1221/MutSigCVsyn)The workflow of MutSigCVsyn is as follows:

The number of synonymous, non-coding, and non-synonymous mutations for each gene (*g*), patient (*p*) and mutation category (*c*) were defined as 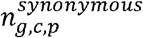, 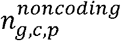 and 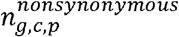. Similarly, the coverage was defined as 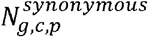, 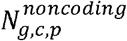, 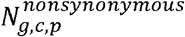. The total count of mutation/coverage across all categories was defined as *c* + 1 as in MutSigCV, whereas for mutations, it meant the sum of all mutations, but in coverage, it meant the sum across categories 1 to 6.

To account for the gene-specific covariates in BMR (background mutation rate), MutSigCVsyn finds the nearest neighbor genes, which share the closest mutational property based on the covariates (expression level, DNA replication timing, and chromatin compartment), for each target gene.

First, as in MutSigCV, the pairwise Euclidean Distance between every gene pair was calculated according to the gene covariate information. For each gene *g*, the raw background mutation number and coverage were defined as the non-coding mutation number and coverage(Eq 1.1) across all patients(*p*)and mutation categories(*c*).

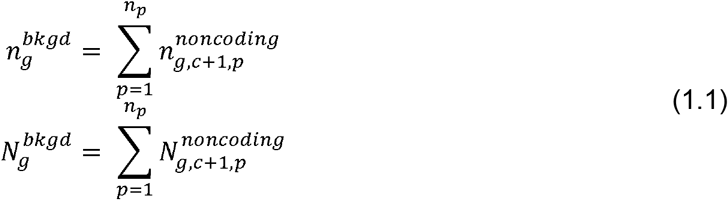

Then, MutSigCVsyn evaluates the non-coding mutation and coverage similarity between pairs of the closest neighbor genes(*i*) and the target genes(*g*) using beta-binomial distribution as in MutSigCV. All qualified neighbor genes (*i* = 0,1,2,…) composed a ‘Bagel’ for the target gene (∀ *i* ∈ *B_g_*). The gene’s background mutations and coverage were calculated by summing the mutation count and coverage (Eq 1.2) across the gene itself and the other qualified genes in its ‘Bagel’.

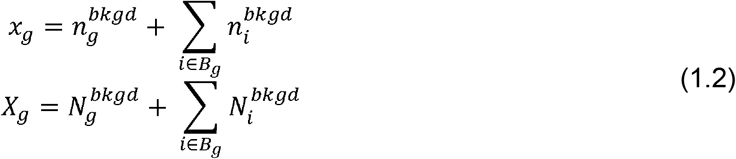

Then, MutSigCVsyn incorporated the marginal relative rate of patient-specific and mutationcategory-specific mutation rate calculated within each histology cohort. The category and patient-specific mutation rate were calculated based upon all mutations (synonymous, non-synonymous, and non-coding) to obtain an accurate estimation of mutational load for each gene. They were then combined with the background mutation count and coverage for the gene of interest to obtain the gene, patient, mutation category level background mutation rate(*x_g,c,p_*) and coverage(*X_g,c,p_*).

After that, for the gene of interest, the probability of observing 0, 1, or more synonymous mutations in each mutation context and patient was calculated (Eq 1.3). Here, the 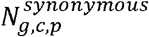 indicates that only the possible mutations that happen in the synonymous positions were considered.

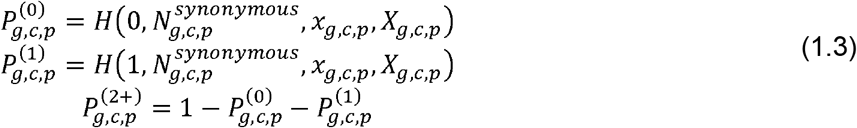

The mutational categories were rank ordered from high to low based on the probability of having 0,1, or more mutations in that category. The probabilities were combined and projected for each 2D combination of the mutation category of the 0, 1^st^, and 2^nd^ mutations and then log-transformed into the scores as in MutSigCV. In addition, the ‘null score boost’, an additional score for deletion and insertion mutations, was set to 0 as synonymous mutations do not fall into this category. A background null distribution was then built by convoluting the mutation probabilities across all 2D projected categories. Finally, the observed score was obtained by summing the scores across observed 2D projected categories of each patient. The p-value for the gene was obtained as the probability of observing a score at least as extreme as the observed score in the null distribution.

The last step was FDR calculation for multiple hypothesis testing. During the identification of genes of which synonymous mutations are significantly mutated, we were identifying signals of substitutions that are commonly known as ‘passenger’ mutations. Therefore, the false discovery rate control would be much more difficult as most of the genes will accept the null hypothesis, leaving a much smaller number of potentially interesting genes for more intensive investigation. Thus, instead of the original Benjamini-Hochberg FDR method, a nonparametric, empirical Bayes FDR method was employed.

### Significant synonymous candidate discovery by Bayesian FDR

The Bayesian false discovery rate as described in Efron et al.^41^ was adopted. Two classes of genes were defined: genes of which the synonymous mutations are significantly mutated, and genes of which the synonymous mutations are not significantly observed. The p-values for each gene are *S*_0_, *S*_1_, *S*_2_,…, *S_N_* to avoid confusion with the probability *p*.

Let the prior probabilities and the hypotheses be:

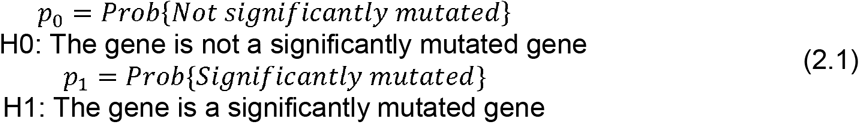

The prior probability has corresponding density *f*_0_(*s*) and *f*_1_(*s*) for the *S_i_* of the gene. Therefore, the mixture density of the 2 populations is

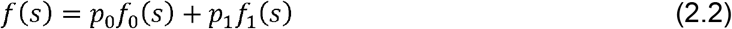

Define *F*_0_(*s*) and *F*(*s*) be the cumulative distribution functions corresponding to *f*_0_(*s*) and *f*(*s*) in (Eq 2.2). According to the definition of Bayesian FDR, The FDR value for {*S* ≤ *s*} is defined as:

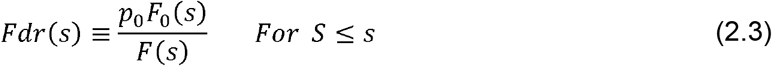

which is the probability of identifying genes coming from the null hypothesis, given p-values equal or less than *s*.

In MutSigCVsyn FDR calculation, a nonparametric estimate for *Fdr*(*s_i_*) was calculated using the empirical CDF of *S*:

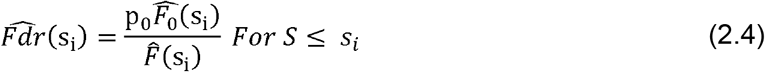

where the *Fdr* value was calculated for every gene *i* with p-value < 0.05.

Note, (1) Both 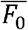 and 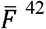 were estimations. To estimate the null distribution, nonexpressed genes (FPKM<1) across all tumor types were used as they are usually regarded to have no role in cancer. 1048 genes in total were used to build the empirical null distribution. The kCDF function in R package sROC^43^ was used for estimating the cumulative distributions. The package gives asymptotically unbiased and consistent estimates for *F*(*s*) and *F*_0_(*s*) given a large number of genes^44^. (2) The conservative assumption that *p*_0_ = 0.99 was adopted because significant candidate genes are expected to occur at a very low chance. (3) As a final step of determining significant candidates, the candidate genes (*i.e*., protocadherin gene families) of which coding and intronic regions are highly clustered in the same genomic regions were excluded^45^ to avoid ambiguity of mutation annotation in overlapped gene regions.

### MutSigCVsyn non-synonymous result analysis

For the drivers in PCAWG, only drivers identified in protein-coding regions were collected (‘element_type’ is ‘cds’). We collected in total 150 PCAWG coding drivers, including drivers discovered previously and drivers discovered exclusively by PCAWG. The 15 PCAWG exclusive drivers were identified by the ‘discovery_unique’ flag in PCAWG.

### Synonymous mutational heterogeneity analysis

The synonymous mutation rate was defined as the rate of synonymous substitutions per 1Mbp synonymous site. The synonymous sites were defined as genome positions where synonymous mutations were likely to occur. For every nucleotide in protein-coding gene sequences, there is 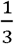 chance for it to mutate into each of the rest nucleotides. Each nucleotide change that caused a synonymous mutation was counted as 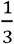 bp. For each patient, the number of total synonymous mutations across all synonymous positions were calculated and the synonymous mutation rate was then calculated as

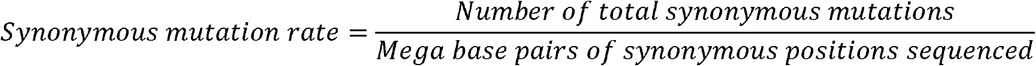

Patients who have 0 synonymous mutations were set to have 0.01 synonymous mutations per mega base pair. The number of synonymous mutations that fell into the mutation category 1-6 was collected and scaled into fractions by the total number of synonymous mutations for each patient

To show local mutation rate variation, chromosome 8 and chromosome 18 were selected and the mutation rate of 3 histology cohorts (Ovary-AdenoCA, Lung-SCC, Thy-AdenoCA) across the entire chromosome were examined. Mutation number in a 1Mbp window sliding over each base pair was collected and averaged across the patient number in that cohort.

### BCL-2 mutation enrichment analysis

The BCL-2 synonymous mutations were extracted from PCAWG Lymph-BNHL maf files. In the permutation analysis, each mutation was randomly assigned to a BCL-2 coding position in one permutation and the number of mutations that fall into the BH4 motif was recorded. After 10,000 permutations, the observed BH4 mutation number and the permuted distribution were compared. The p-value is calculated as

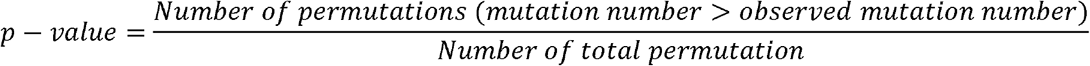

### Gene mRNA expression analysis and CERES score analysis

Gene mRNA expression data in patient tumor sample and normal sample(if exists) were collected. For DepMap cell line expression analysis, the cell line lineage that matches the corresponding histology cohort was first retrieved. The expression of the gene in the cell line lineage was then extracted and compared to all other cell lines. The Mann-Whitney U test was then performed to determine the significance of the difference in gene mRNA expression.

## Results

### Synonymous mutation rate varies across patients, tumor types and genes, impeding driver discovery

The accurate identification of non-synonymous drivers requires explicit corrections for background mutation biases across patients, genes, and diseases. We first examined if the same should be done when identifying synonymous drivers because it is highly likely that there are distinct synonymous mutation rates across these categories.

To demonstrate this synonymous mutation heterogeneity, we collected synonymous mutations in 18638 protein-coding genes across 2572 white-list PCAWG patients. We calculated the rate of synonymous substitutions per 1Mbp synonymous site for each indication (see Methods Section). As expected, the synonymous mutation rate was lower than the total mutation rate(Figure 1a, top). We observed that the synonymous mutation frequencies vary widely across patients and histology indications. Across the indications, Skin-Melanoma has the highest median synonymous mutation frequencies across patients at 21.7 per Mbp. Towards the other extreme, the lowest median frequency is observed in CNS-PiloAstro(0 per Mbp, due to patients having no synonymous mutations) which is over 20 times smaller than Skin-Melanoma.

**Figure 1.**
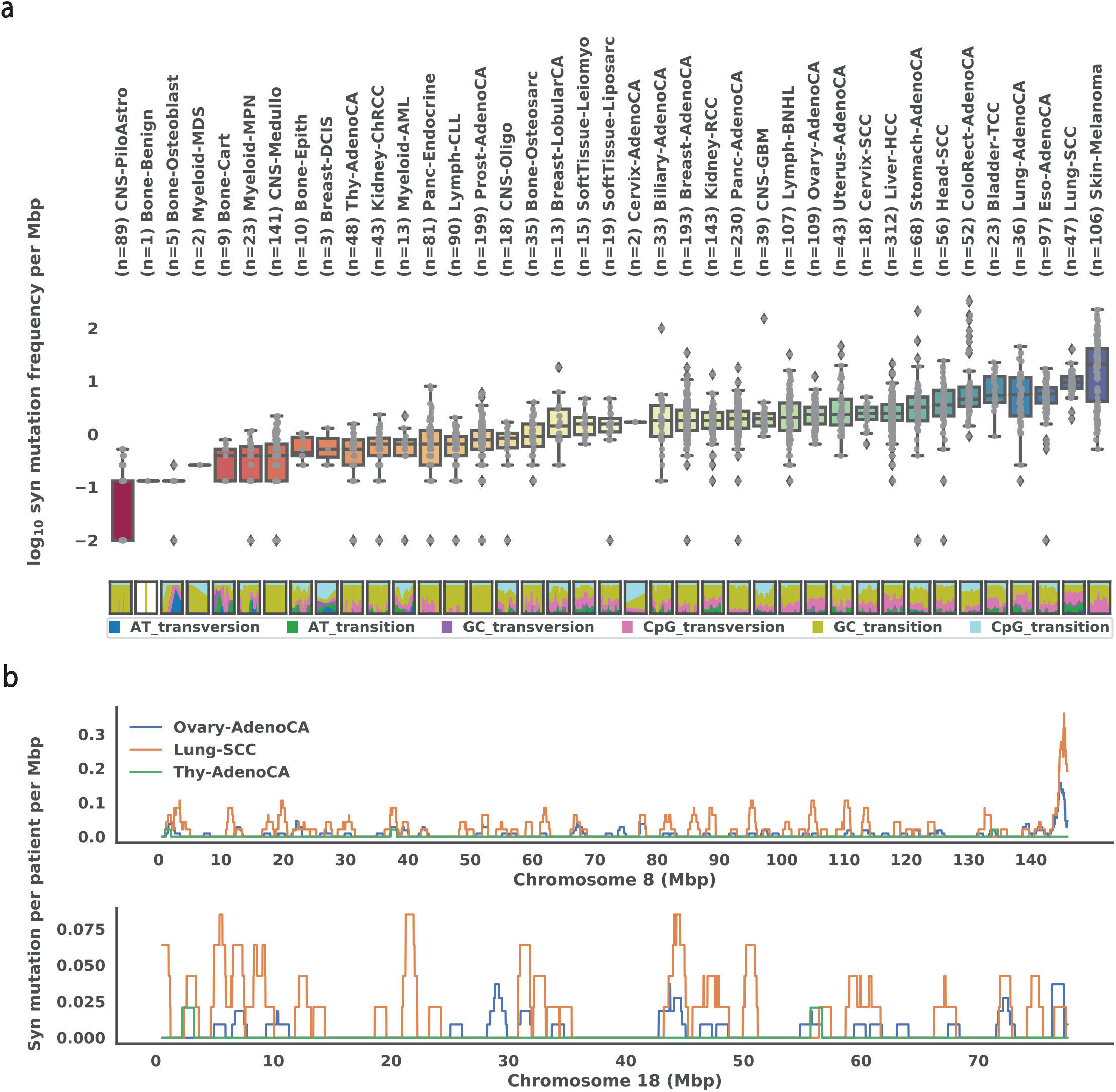
Synonymous mutation rate in cancer varies across patients, histology types, and genes. (A) Box plot (Top) of patient synonymous mutation frequency across all histology types. Mutation frequency is shown as logarithmically transformed mutation number per mega basepair. Patients that don’t have any synonymous mutations are set to have −2 transformed mutation frequency per mega basepair. Each dot represents a patient. Histology types are ordered by their median somatic mutation frequency. The relative percentage(bottom) of mutations falls into 6 mutation categories(see Methods Section) for all individual patients across the histology types. (B) Synonymous mutation number averaged by patient number in Ovary-AdenoCA (blue), Lung-SCC (orange), and Thy-AdenoCA (green), respectively, illustrated on the entire chromosome 8 (top) and chromosome 18 (bottom).

We also observe large variations in mutation frequency within individual cancer indications. Except for some of the extremely small cohorts (e.g., Bone-Benign (n=1), Bone-Osteoblast (n=5), Myeloid-MDS (n=2), Cervix-AdenoCA (n=2)), the maximum mutation frequency is at least 1 order of magnitude larger than the minimum in each indication. The largest such variation occurs in ColoRect-AdenoCA, where the highest synonymous mutation frequency is 329 per Mbp, while the lowest is 0.917 per Mbp. This is consistent with the existence of a hypermutated microsatellite instability subpopulation^46^.

These variations are partly explained by mutational etiology (Figure 1a, bottom). A typical example is Skin-Melanoma, which exhibits an enrichment of GC transition mutations, consistent with the known mutational signature due to UV radiation^47^. In addition, the high content of GC transition in Bladder-TCC patients is likely caused by APOBEC protein family activity, which is a prominent mutational signature pattern in TCGA bladder tumors^48^. In Lung-SCC, we also observe signs of signatures related to tobacco smoke, which is characterized by G to T transversion caused by lesions when polycyclic aromatic hydrocarbons enter the human body^49^. Thus, as expected, known mutational signatures contribute to synonymous mutation heterogeneity as well.

Next, in order to illustrate the heterogeneity of mutation rate across genomes for a given cancer indication, we plotted the average synonymous mutation number per patient across chromosome 8 and chromosome 18 for 3 histology cohorts(Ovary-AdenoCA, Lung-SCC, Thy-AdenoCA). As shown in Figure 1b, variation of local mutation numbers is observed across all 3 histology types.

These results demonstrate that there is substantial variability in the synonymous mutation burden at the histology, patient, and gene levels. Therefore, the assumption of a constant mutation rate and completely independent mutation events is not appropriate for synonymous driver identification. To accurately identify synonymous drivers, driver predictions must explicitly correct for these covariates.

### MutSigCVsyn detects differences between observed and expected synonymous mutation frequencies in cancer cohorts

In order to correct for these covariates, especially the gene-specific differences in mutation rate, we adopted and modified MutSigCV^4^ (Figure 2a), which corrects for variation by using patientspecific mutation frequencies and the 192-triplet nucleotide mutation context (e.g., A(A->C)A), and gene-specific background mutation rates through the incorporation of expression level and chromosome replication position.

**Figure 2.**
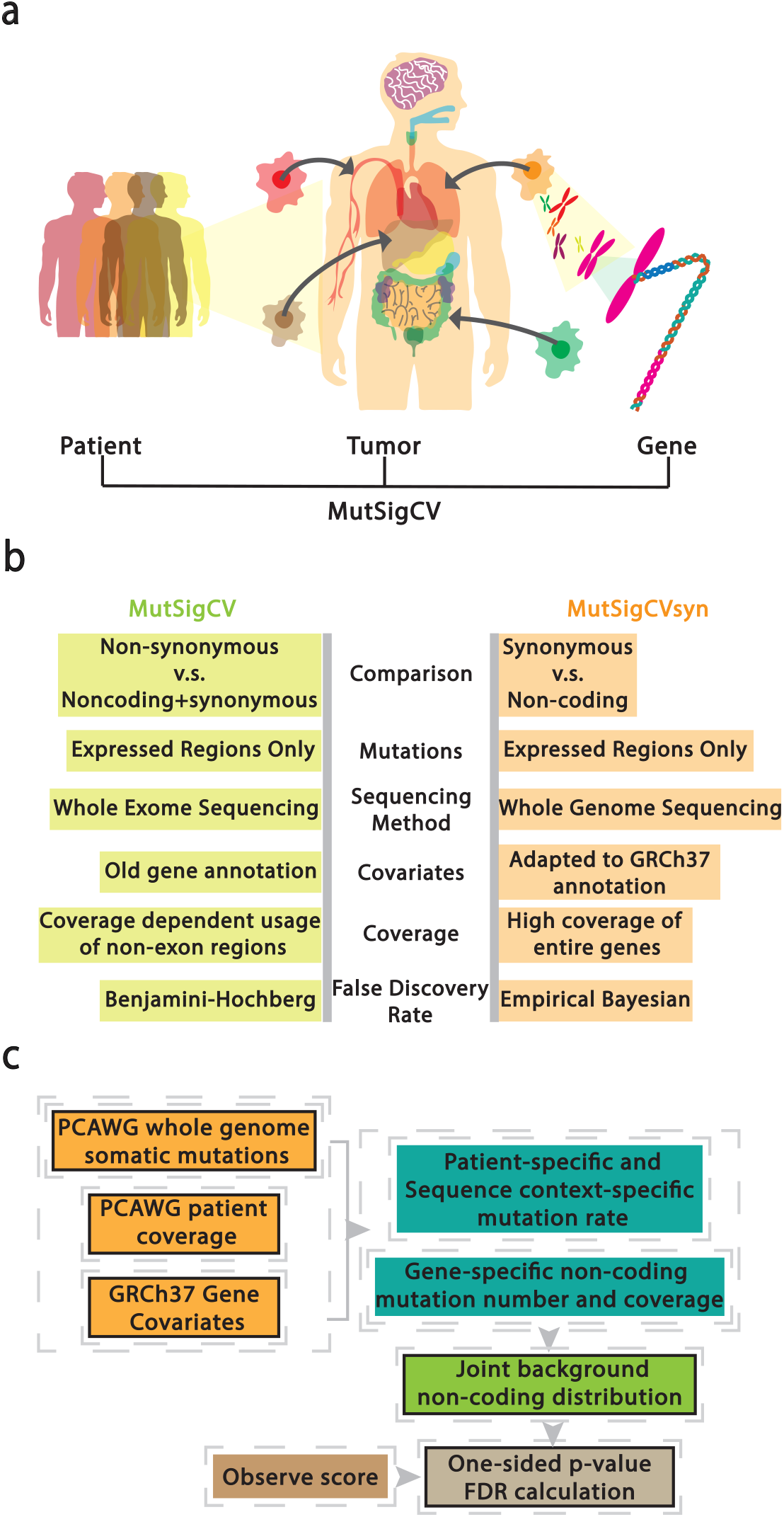
Changing MutSigCV to MutSigCVsyn to identify synonymous cancer drivers. (A) MutSigCV accounts for mutation heterogeneity across patients, diseases, and genes. (B) Comparison between MutSigCV and MutSigCVsyn: (1) MutSigCVsyn uses only non-coding mutations instead of background comprised of both non-coding and synonymous mutations adopted by a majority of driver mutation detection algorithms. (2) MutSigCVsyn utilizes whole genome sequencing input data instead of whole-exome sequencing. (3) Both MutSigCVsyn and MutSigCV only utilize mutations in transcriptionally expressed regions. (4) MutSigCVsyn utilizes a re-annotated covariate file that was adapted to the PCAWG Gencode v19 annotation. (5) MutSigCVsyn patients have high-quality coverage data over non-coding regions, compared to limited coverage in the original MutsigCV. (6) MutSigCVsyn utilizes a non-parametric empirical Bayesian method to calculate local FDR value. (C) The outline of MutSigCVsyn. Boxes with solid lines show MutSigCVsyn exclusive input/steps (see Methods section for detailed description).

MutSigCV was originally designed for the identification of non-synonymous drivers in the context of exome sequencing data. To convert MutSigCV into a synonymous driver detection algorithm, we made several modifications (Figure 2b). The biggest modification is using only the non-coding mutations in our background mutation model. The original MutSigCV’s background model is composed of synonymous mutations and non-coding mutations found in the untranslated regions of transcripts, but with limited coverage in non-coding regions. This is because it was originally designed for cancer exome re-sequencing datasets. However, the high data quality and coverage in PCAWG Whole Genome sequencing datasets allow us to use the mutations in the complete intronic region and untranslated regions for the mutational background The 2 major reasons for using such a background are: (1) we adopted a simplifying assumption that on average, non-coding mutations are ‘more neutral’ than the synonymous mutations. The lower rate in the intronic region than in exonic regions across species^50^ suggests non-coding regions of genes are under weaker selection than the coding region. (2) By restricting the non-coding mutations to the mutations occurring in transcribed regions, we prevent bias caused by different mutation frequencies in transcribed versus non-transcribed regions. Specifically, the non-coding mutations in our analysis only include (a) intronic mutations and (b) mutations in untranslated regions.

In MutSigCVsyn, protein-coding gene coverage information for every patient in PCAWG is calculated. In addition, we re-annotated the gene covariate file to adapt the gene name annotation in PCAWG. To benefit the community, files and scripts are available publicly on GitHub. The workflow of MutSigCVsyn is shown in Figure 2c. A more detailed description of MutSigCVsyn can be found in the Methods Section.

### Quality control: MutSigCVsyn identifies non-synonymous drivers with high sensitivity

MutSigCVsyn is designed for the identification of synonymous drivers. However, if MutSigCVsyn builds a valid non-coding background, MutSigCVsyn should be able to identify non-synonymous drivers as well. Therefore, as quality control for our approach, we applied MutSigCVsyn to 2572 donors in 39 PCAWG histology types to identify non-synonymous drivers, using non-coding mutations as background.

We identified a total of 133 significant genes (Figure S1) across 29 cohorts. As expected, most of the genes in the candidate gene list have been reported before. As the most frequently altered gene in human cancer, TP53 is the most frequently significant driver across all indications. It is called significant in 21 out of 39 histology types, including ColoRect-AdenoCA, Lymph-BNHL, Liver-HCC, and Panc-AdenoCA. Furthermore, our significant driver list for each indication overlaps the known cancer drivers in that indication. We identified candidate genes CDKN2A in HCSCC (Head and Neck Cancer), which is a known tumor suppressor and whose inactivation has been well studied in HCSCC^51^. In CRC (Colorectal Cancer), APC and SMAD4 are also identified as the candidates. APC constitutively activates the canonical WNT signaling in most colorectal cancer cases, leading to cell proliferation and tumor formation^52^. Another known gene, SMAD4^53^, which negatively regulates TGF-beta, is also frequently found in CRC patients. Finally, our results in Breast-AdenoCA also highlighted some genes that are specifically known to be frequently mutated in breast cancer^54^, including PIK3CA, CDH1, GATA3, MAP2K4.

As a further test of our result, we compared our output to CGC (Cancer Gene Census) and PCAWG driver list (see Methods Section) (Figure 3a). We observed 58.6% (78 out of 133) of our non-synonymous list overlaps with the CGC genes. The high overlap rate may be due to the nearly full coverage of non-coding regions and the accurate calculation of the coverage file for the analysis. We also observe 66.2%(88 out of 133) of our candidate genes overlap with PCAWG drivers. Additionally, we successfully identified 6 genes out of the 15 PCAWG exclusive drivers (Table S1), which are the genes identified in the PCAWG cohort for the first time. In conclusion, these results indicate that our modifications to MutSigCV do not dramatically affect the ability of MutSigCVsyn to reproduce previously known results.

**Figure 3.**
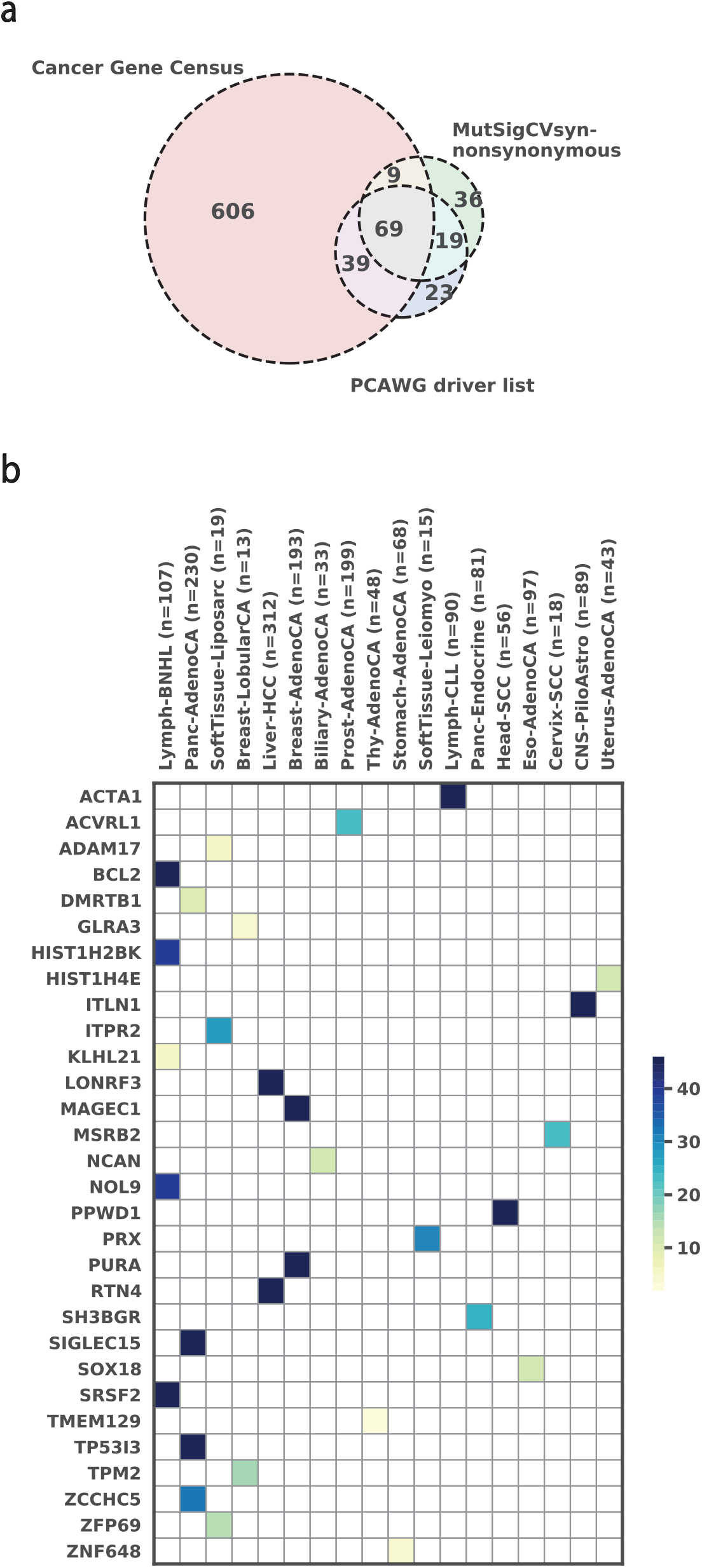
MutSigCVsyn identifies non-synonymous and synonymous cancer drivers. (A) Venn Diagram displaying overlapped gene numbers of MutSigCVsyn significant non-synonymous drivers with Cancer Gene Census and PCAWG driver lists. (B) Heatmap shows significant synonymous candidate genes (Bayesian FDR < 1× 10^(−2)) identified by MutSigCVsyn. Genes are colored by the negative logarithm of the transformed FDR value from high (dark blue) to low (light yellow).

### The landscape of synonymous drivers

Given our ability to identify non-synonymous drivers with high sensitivity, we used MutSigCVsyn to identify synonymous drivers in all 39 histology types in PCAWG. We identified 30 significant synonymous candidates in total (Figure 3b). As expected, this list is parsimonious and smaller than the non-synonymous driver list. Lymph-BNHL has the most significant synonymous candidates (n=5), followed by Panc-AdenoCA(n=4). In total, there are 18 distinct indications having significant genes. The variety of indications implies that MutSigCVsyn is not biased by histology-wise mutation frequencies. Among all candidates, 11 genes across 7 indications have the smallest p-values (p-value < 1.0 × 10^−7^), including BCL2 and SRSF2 (Lymph-BNHL), ITLN1 (CNS-PiloAstro), PPWD2 (Head-SCC), PURA and MAGEC1 (Breast-AdenoCA), SIGLEC15 and TP53I3 (Panc-AdenoCA), etc.

Two of the top candidates, BCL-2 and SRSF2, are known to be non-synonymous drivers of cancer as cataloged in the Cancer Gene Census. Both genes were identified in the Lymph-BNHL cohort. The t(14;18) translocation in BCL-2 is critical in follicular lymphoma progression^55^ and SRSF2 is a global splicing regulator that binds to exonic splicing motifs. It is associated with hematopoietic diseases (i.e., myelodysplastic syndrome^56^), but hadn’t been specifically characterized in Non-Hodgkin Lymphoma. In PCAWG, 3 unique Lymph-BNHL patients have 3 distinct synonymous mutations in SRSF2: p.Y3Y (DO27764), p.V79V (DO52664), and p.G82G (DO52672), the latter 2 reside in the RNA recognition motif (RRM)of SRSF2. Though one of the patients (DO52664) carried missense mutation at Proline95 position that is known to alter mRNA binding affinity^57^, 2 other patients only harbor SRSF2 synonymous mutations. As synonymous mutations can disturb mRNA translation initiation and elongation process^58^, it is possible that SRSF2 synonymous mutations alter RRM binding affinity and contribute to a global transcriptional profile change in cancer cells.

While many of the candidate genes are poorly studied in cancer, there is evidence to suggest some of them could be required for tumor growth. For example, PURA, which encodes the nucleic acid-binding proteins Pura, is one of the significant candidates in Breast-AdenoCA. Studies have found that overexpression of PURA inhibits proliferation and anchorage-independent colony formation of Ras-transformed NIH3T3 Fibroblast cells, suggesting PURA acts as a potential tumor suppressor gene^59^. In our analysis, the PURA expression level is significantly lower(Mann-Whitney U-test p-value = 5 × 10^−3^) in tumor samples(n = 85) than in normal samples(n = 6) (Figure S2a). This low expression suggests a plausible contribution to breast cancer cell proliferation. Another example is the immune checkpoint gene SIGLEC15, the top significant gene in Panc-AdenoCA. SIGLEC15 is a well-conserved member of the immunoglobulin superfamily of receptors Siglecs that bind to sialic acid. In a recent study^60^, upregulated SIGLEC15 has been widely found across different cancer types and had been related to a worse patient survival rate. Moreover, SIGLEC15, rather than other immune checkpoint genes, was found to have a positive expression correlation with upregulated genes in pancreatic cancer^61^. We observe a significantly higher expression of SIGLEC15 mRNA expression in Pancreas exocrine lineage cancer cell lines than in other cell lines in DepMap (Mann-Whitney U-test p-value = 5 × 10^−3^)(Figure S2b). We used DepMap data because the transcriptome data of PCAWG pancreatic patient normal specimens is unavailable. Combining with SIGLEC15’s mutually exclusive expression with B7-H1(PD-L1)^62^, we think that SIGLEC15 levels may play a role in pancreatic cancer immune evasion. The role of synonymous mutations in both cases is therefore worthy of further study.

### MutSigCVsyn exclusive synonymous candidates might contribute to cancer

We expect that the significant synonymous drivers called by MutSigCVsyn have the potential to contribute to a cancer phenotype. We focus on one of our particular candidates, BCL-2, that has compelling cancer associations. BCL-2(B cell lymphoma 2) regulates apoptosis by antagonizing the action of proapoptotic BCL-2 family members^63^. It was originally identified as the protooncogene involved in the t(14;18) translocation in follicular lymphoma^64^. Among the BCL-2 protein motifs, the BH4 motif is essential for the anti-apoptotic activity of BCL-2. The deletion of the BH4 region in a human fibroblast cell line largely impairs the cell viability under IL-3 deprivation^65^ and melanoma growth in vitro and in vivo^66^.

In our analysis, we observe 41 synonymous mutations in 26 unique patients, and 9 of the mutations in 9 different patients reside in the BH4 motif (Figure 4a). Combining with the anti-apoptotic effect of the BH4 motif, we hypothesize that there may be an enrichment for synonymous mutations in the BH4 motif that might disrupt its function and thus promote cancer cell survival. If this is true, we would expect to see a significant enrichment of synonymous mutations in the BH4 motif vs the BH1,2, or 3 motifs. To test for enrichment, we conducted a permutation test for the observed number of mutations in the BH4 motif by comparing it to the 10000 permutations where all 39 mutations are randomly assigned across all the BCL-2 coding positions. The results show that BCL-2 synonymous mutations are significantly enriched in the BH4 motif (p-value = 0.032) (Figure 4b). This indicates that synonymous mutations in the BCL-2 BH4 motif might be positively selected in lymphomas.

**Figure 4.**
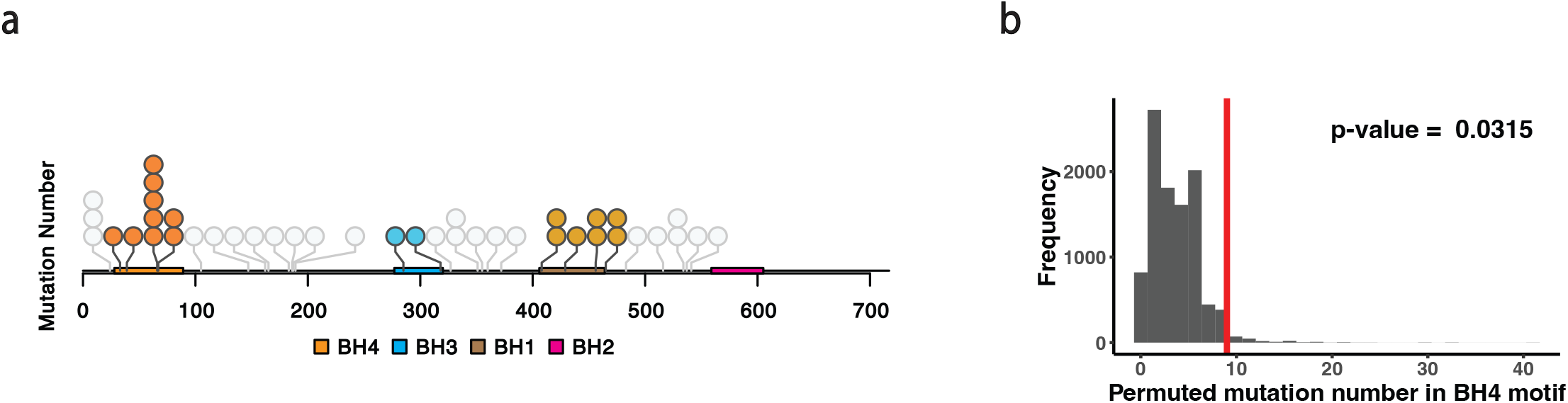
Synonymous mutations are significantly enriched in BCL-2’s BH4 motif. (A) Illustration and distribution of all BCL-2 synonymous mutations identified in PCAWG Lymph-BNHL patients across the BCL-2 coding sequence. Dots represent the occurrence of each synonymous mutation. BCL-2 motifs and the synonymous mutation in those motifs are colored: BH4 (Orange), BH3 (Blue), BH1 (Brown), BH2 (Bright Pint). Synonymous mutations that fall outside of the motifs are colored grey. (B) BH4 synonymous mutation number distribution from permutation test (permutation number = 10000). The red line shows the observed number of synonymous mutations (n=9).

## Discussion

MutSigCVsyn controls for patient-, histology-, and gene-specific mutation rate variations to identify synonymous cancer drivers. What is novel and significant about this approach is that the background mutation model we constructed accounts for the covariates that are the gold standard for properly identifying non-synonymous drivers. Without adjusting for these covariates, there is a high likelihood of misidentifying synonymous cancer drivers. To test this approach, we reasoned that our new background mutation model should still be able to identify known non-synonymous drivers. And indeed, we find MutSigCVsyn identifies above 60% of the drivers reported in the CGC. 60% is a high success rate, given that an evaluation of eight different driver-gene-detection algorithms^67^ found that they identified between ~ 10% and 50% of the drivers in CGC.

By applying MutSigCVsyn to the PCAWG database, we identified 30 synonymous cancer drivers. Among them, BCL-2 appears to be the most promising candidate due to the extensive literature concerning its role in follicular lymphoma^55,68,69^ and the significant clustering of synonymous mutations in BCL-2’s BH4 regulatory motif. Thus, we hypothesize that synonymous drivers in the BH4 motif might contribute to BCL-2’s gain-of-function role in oncogenesis. One potential argument against this hypothesis is that the enrichment of mutations in BCL-2 is the result of somatic hypermutation caused by activation-induced cytidine deaminase, which is frequent in immunoglobulin variable regions^70^. However, we find that only 4 of the 26 patients that have synonymous mutations in BCL-2 harbor an IgG translocation in this gene. Further, the breakpoints of the BCL2 translocation within these 4 patients are at least 100kbp away from the observed synonymous mutations - a distance that is not consistent with hypermutations in immunoglobulin variable regions. And most importantly, the background estimate of the activation-induced-cytidine-deaminase signature is already accounted for in the MutSigCVsyn analysis, as well as other candidate hypermutations in lymphomas, meaning they are statistically excluded from our candidate list. Thus, these results suggest that synonymous mutations could result in a gain-of-function in BCL-2, which might be positively selected for in lymphoma patients.

The divergence at nonsynonymous and synonymous sites in cancer cohorts, known as the dN/dS ratio, is a conventional measure of evolutionary selection pressure^71^. It has been applied in many somatic evolution studies^72–75^ under the assumption that nearly all synonymous mutations are neutral^14^. A small dN/dS ratio is usually interpreted as a global signal of negative selection on non-synonymous mutations. BCL-2 challenges this interpretation. In a study by Lohr et al.^76^, a small dN/dS ratio was found across the entire BCL-2 gene in a 50 diffuse large B-Cell lymphoma patient cohort. It was thus concluded that BCL-2 undergoes strong negative selection. Contradictory to this, our study suggests that an increase in dS creates a robust positive selection signal of synonymous mutations in the BH4 motif of BCL-2. Thus, it may not entirely be that evolution is selecting negatively on the numerator dN, but rather, positively on the denominator dS. Therefore, the possibility exists that the negative selection pressures on BCL-2 are overestimated when only using the dN/dS ratio across the entire gene. More broadly, this indicates that the interpretation of the dN/dS ratio may not be straightforward when synonymous mutations are not neutral.

Except for BCL-2^34^, the other candidate genes identified by MutSigCVsyn (Figure 3b) have not been identified previously. Differences in datasets and methodology are two reasons differences in the published lists of synonymous cancer drivers can arise. For example, PCAWG, which we used in this study, is less comprehensive than COSMIC in terms of the number and source of identified synonymous mutations. However, PCAWG uses uniform analysis standards, whereas COSMIC uses human curation of publications reporting somatic mutation results based on heterogeneous analysis standards (e.g., differences in alignments, variant callers, manual annotations) – which may affect accuracy.

By not accounting for covariates in a background mutation model, studies can identify spurious synonymous mutation candidates. In one study^35^, multiple mutations in two extremely long human genes, which encode the muscle protein titin and neuronal synaptic vesicle protein piccolo, were identified as synonymous cancer drivers. However, these genes are commonly observed false positives in non-synonymous driver identification studies that don’t account for the gene-specific mutational biases^4^. In another study^34^, two top synonymous candidates are present in highly mutable microsatellite regions: MLLT3 (c.501T>C) has the 5^th^ highest SynMICdb score, and ARID1B (c.768C>A) has the 13^th^ highest SynMICdb score. Creating an appropriate background mutation model minimizes such microsatellite biases. Therefore, these putative false-positive results highlight the importance of methodologies that utilize comprehensive background mutation models, of the type adopted by MutSigCVsyn.

To establish a mutational background with which to compare synonymous mutations, MutSigCVsyn utilizes the non-coding mutations in the UTR and intronic regions. There are potential drawbacks to utilizing this non-coding background. Sequences in some non-coding regions are under evolutionary constraints, especially regulatory elements, such as at intron-exon junctions^77,78^. A positively selected non-coding background may diminish the synonymous mutation signal and decrease the number of synonymous drivers. For these reasons, it may be useful in the future to exclude specific background regions that are already known to be under evolutionary selection. However, principled exclusion criteria will require much larger cohorts and more complete knowledge of positive selection in non-coding regions of the genome.

In conclusion, MutSigCVsyn is the first synonymous cancer driver identification algorithm that uses the standards that are commonly found in algorithms for non-synonymous cancer drivers. We have identified a list of 30 novel synonymous drivers that provide promising opportunities for future experimental research to understand how these synonymous drivers can contribute to cancer.

## Acknowledgement

JP acknowledges funding from NIBIB (5R21EB026617). EPO acknowledges funding from NIH (R35-GM124818) and NSF (ABI-1759860).

---

**Figure S1.**
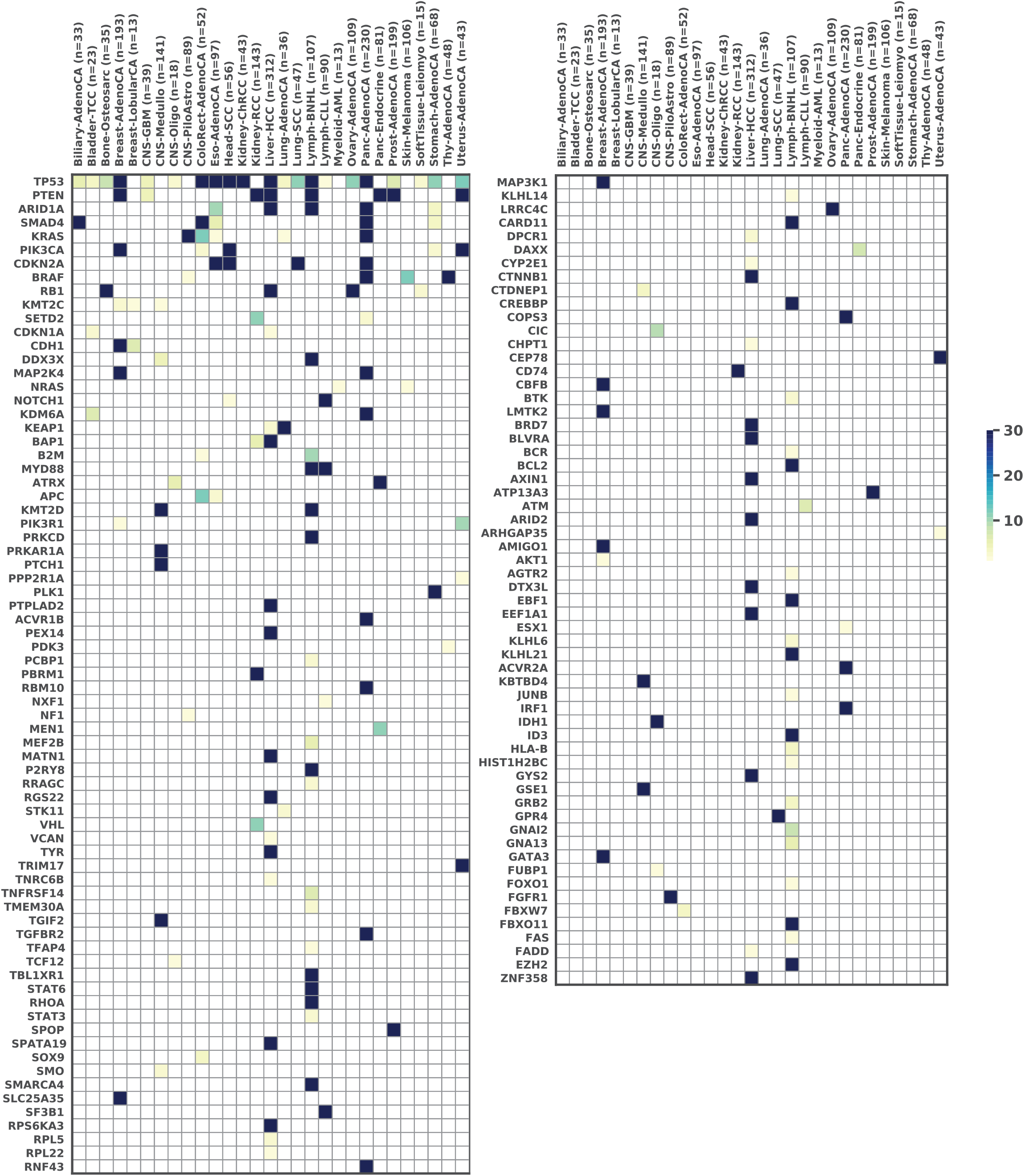
MutSigCVsyn non-synonymous cancer driver landscape. Heatmap displaying 133 significant non-synonymous candidate genes (Benjamini-Hochberg FDR < 1× 10^(−2)) identified by MutSigCVsyn. Candidate genes are divided into two columns and are ranked from most frequently across histology cohort (left top) to the least frequent ones (right bottom). Candidate genes are colored by negative logarithmic transformed FDR value from high (dark blue) to low (light yellow).

**Figure S2.**
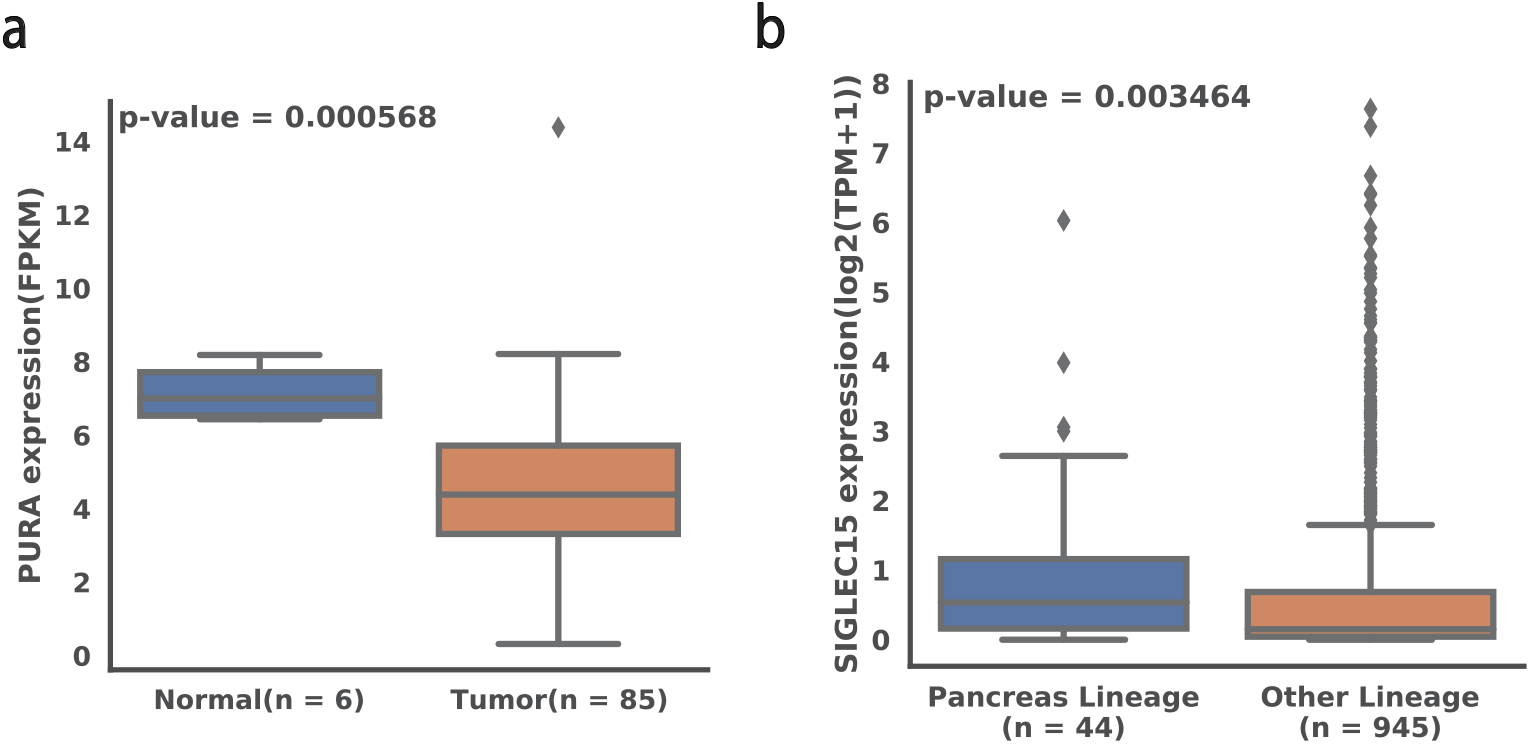
Potential functional role of MutSigCVsyn synonymous drivers. (A) Boxplot of Breast-AdenoCA patient PURA mRNA expression level of normal samples and tumor samples. P-value is calculated by the Mann-Whitney U test. (B) Boxplot of SIGLEC15 expression data from DepMap Pancreas exocrine cell lines and all other tested cell lines. P-value is calculated by the Mann-Whitney U test.

**Table S1.** MutSigCVsyn identifies PCAWG exclusive drivers in non-synonymous analysis. PCAWG exclusive drivers are the cancer driver genes that were first identified by the PCAWG working group. There are in total 15 exclusive non-synonymous protein-coding drivers in PCAWG and they are shown in the table. The ‘gene’ column shows the gene name. ‘cds’ in the Element_type’ column shows that the coding region of the gene is identified as a cancer driver. ‘discovery_unique’ in the ‘category’ column shows that the gene is first identified by PCAWG. 6 of them (highlighted yellow) were identified by MutSigCVsyn in non-synonymous mutation analysis.

| Gene | Element_type | Category |
| --- | --- | --- |
| TMEM30A | cds | discovery_unique |
| PLK1 | cds | discovery_unique |
| PA2G4 | cds | discovery_unique |
| SRSF7 | cds | discovery_unique |
| CAMK1 | cds | discovery_unique |
| TMSB4X | cds | discovery_unique |
| KLHL6 | cds | discovery_unique |
| RRAGC | cds | discovery_unique |
| GRB2 | cds | discovery_unique |
| DYRK1A | cds | discovery_unique |
| CTC-512J12.6 | cds | discovery_unique |
| DYNC1I1 | cds | discovery_unique |
| PRKCD | cds | discovery_unique |
| RIPK4 | cds | discovery_unique |
| RELA | cds | discovery_unique |

## Notes

### Competing Interest Statement

JRP has ownership interests in Theseus Pharmaceutical and MOMA therapeutics.
JRP is a co-founder of Theseus pharmaceuticals.
JRP is a paid consultant for Takeda Pharmaceuticals, Theseus Pharmaceuticals, MOMA therapeutics, and third rock ventures.

